# Multi-omics computational analysis unveils the involvement of AP-1 and CTCF in hysteresis of chromatin states during macrophage polarization

**DOI:** 10.1101/2023.09.27.559849

**Authors:** Yubo Zhang, Wenbo Yang, Yutaro Kumagai, Martin Loza, Sung-Joon Park, Kenta Nakai

## Abstract

Macrophages display extreme plasticity, and the mechanisms and applications of polarization and de/repolarization of macrophages have been extensively investigated. However, the regulation of macrophage hysteresis de/repolarization remains unclear. In this study, by using a large-scale computational analysis of macrophage omics data, we report a list of hysteresis genes that maintain their expression patterns after polarization after ulti- and de/repolarization. While the polarization in M1 macrophages leads to a higher level of hysteresis in genes associated with cell cycle progression, cell migration, and enhancement of the immune response, we found weak levels of hysteresis after M2 polarization. During the polarization process from M0 to M1 and back to M0, the factors IRFs/STAT, AP-1, and CTCF regulate hysteresis by altering their binding sites to the chromatin. Overall, our results show that a history of polarization can lead to hysteresis in gene expression and chromatin changes over a given period. This study contributes to the understanding of de-/repolarization memory in macrophages.

## Introduction

Macrophages, an essential cell type of the innate immune system, migrate into different tissues from the peripheral blood and undergo differentiation upon receiving various immune cues (1). The major function of macrophages is to bridge the innate immune system with the adaptive immune system through several biological processes, such as phagocytosis of pathogens/cell debris, antigen presentation on major histocompatibility complexes, and production of cytokines (1–3). Non-activated M0 macrophages polarize into either the classical pro-inflammatory M1 phenotype or alternative M2 phenotype. M1 is typically induced by IFN-γ or Toll-like receptor (TLR) ligands and produces redox molecules (4). M2 macrophages are induced by an exposure to a set of cytokines, and are observed in conditions, such as helminth infection (5), wound healing, cancer, and obesity (6). The polarized macrophages express pro-inflammatory cytokine genes, including IL-1β, TNF, IL-12, and IL-18 in M1; and Chi3l1, Ym1, Rentla, and Arg1 in M2 (7).

Interestingly, macrophages can retain their plasticity. This means that M1 cells exposed to M2-inducing signals can differentiate into M2 states and vice versa, which often influences the outcomes in therapeutic treatment of diseases, such as cancers (8). The transcriptional phenotypes in the cellular microenvironment and subsequent immune response are influenced by not only the ongoing de-/repolarization (9), but also the memory of the past cellular state, which is referred to as hysteresis (10,11). For instance, trained immunity is a typical phenomenon that may be attributed to hysteresis (12), and recent experimental evidence supports the significant influence of hysteresis on immune responses (11).

However, the molecular mechanisms underlying mmacrophage plasticity and cell polarization-mediated hysteresis remain unclear. Therefore, in this study, we conduucted a large-scale computational analysis to investigatee the regulatory mechanisms of macrophage hysteresis duuring polarization. Consequently, we identified hysteresis-related genes that are enriched among interfeeron-stimulated genes (ISGs) and involved in fundamental immune functions. Remarkably, the hysteresis induced byy M1 stimulation was more pronounced than that induced by M2 stimulation. We demonstrated that specific DDNA-binding proteins are tightly associated with different hysteresis strengths. AP-1 and CTCF were consistentlyy enriched in accessible and inaccessible distal regions, resspectively, during M1 hysteresis. These findings suggesst an impact of chromatin remodeling on the regulation of hysteresis genes.

## Results

### Differential gene expression analysis reveals the existence of hysteresis genes after de-/repolarization

We used the time-course RNA-seq data (Figure 1A)) obtained from the Liu project to investigate the influence of de-/repolarization on the macrophage transcriptome. This dataset employed M0 cells as the starting point (0 h)), followed by stimulating the cells with either M1 or M2 cues for 24 h to yield M1 and M2 cells (24 h). After this 24-hour stimulation, the cells were either left untreated for 966 h to facilitate de-polarization (de-polarization group) or subjected to the opposite stimulus for 96 h to facilitate repolarization (re-polarization group) (Figure 1B). Thus, we established the following four cellular states: de-polarized M0 from M1 (deM0_M1), de-polarized M0 from M2 (deM0_M2), re-polarized M1 from M2 (reM1_M2), and repolarized M2 from M1 (reM2_M1). The expression alterations observed in the transcriptome due to de-/repolarization within the same macrophage phenotype were collectively defined as hysteresis. Genes exhibiting significant expression discrepancies were identified as hysteresis genes, which likely mirror the past state of the cells.

We compared the expression profiles of primary macrophages (M0, M1, and M2) with those of the corresponding de-/repolarized macrophages. To elucidate the effects on the transcriptional profile of distinct polarization histories, we compared the primary M0 cells with deM0_M1 and deM0_M2 conditions. By taking into account an absolute log_10_(FPKM) change > 0.5 and P-value threshold of 0.05, 75 (Supplementary Excel Sheet 1) and 31(Supplementary Excel Sheet 2) genes were defined as significant hysteresis genes for the M0 vs deM0_M1 and M0 VS deM0_M2 group, respectively (Figure 2A). Within these two sets of significant hysteresis genes, 24 common genes, including *S100a8, S100a9, Retnlg, Plac8, Ccr2*, and *Bgn*, were identified. In addition, 51 genes, represented by *Ccl5, Irf7*, and *Ccr3*, displayed significant differential expression in the deM0_M1 group compared to that in the M0 group, but not in the deM0_M2 group, thereby highlighting the existence of a stronger M1 hysteresis. In contrast, seven hysteresis genes, including Retnla, were found to be present only in the deM0_M2 group.

**Figure 1:**
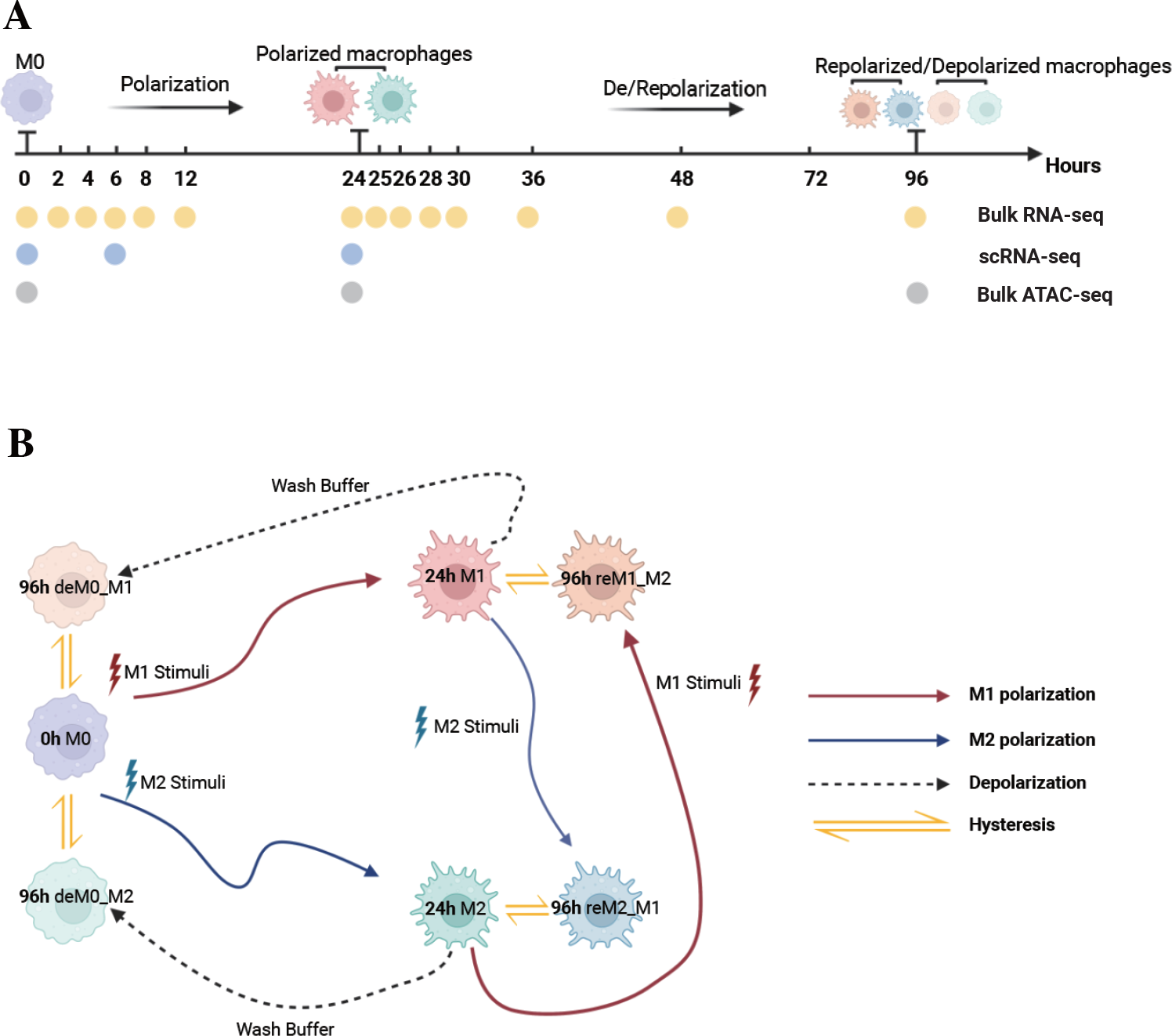
Multi-omics data used for macrophage analysis and explanation of macrophage hysteresis. **(A)** Schematic illustration depicting the macrophage response to cytokine treatment and acquisition of multi-omics data. The horizontal time axis signifies distinct time points (in hours) and the arrows symbolize various types of stimuli. The labels 0 h, 24 h, and 96 h correspond to the pivotal macrophage states. Positioned beneath the time axis, the dots and the accompanying text indicate instances of data collection and respective acquired data types, respectively. **(B)** Schematic representation of macrophage polarization in the context of the original research and delineation of macrophage hysteresis. Each distinct pattern displayed in the colored cells symbolizes the cellular state of macrophages. The colored trajectories and arrows denote the various stimuli (M1 stimulus, M2 stimulus, and wash buffer) and polarization trajectories encountered by the macrophages. The double-headed arrow signifies the variation in expression between the macrophage post-depolarization/repolarization and baseline macrophage states, encapsulating the concept of macrophage hysteresis. (deM0_M1: depolarize M0 from M1, deM0_M2: depolarize M0 from M2, reM1_M2: repolarize M1 from M2, and reM2_M1: repolarize M2 from M1.)

**Figure 2:**
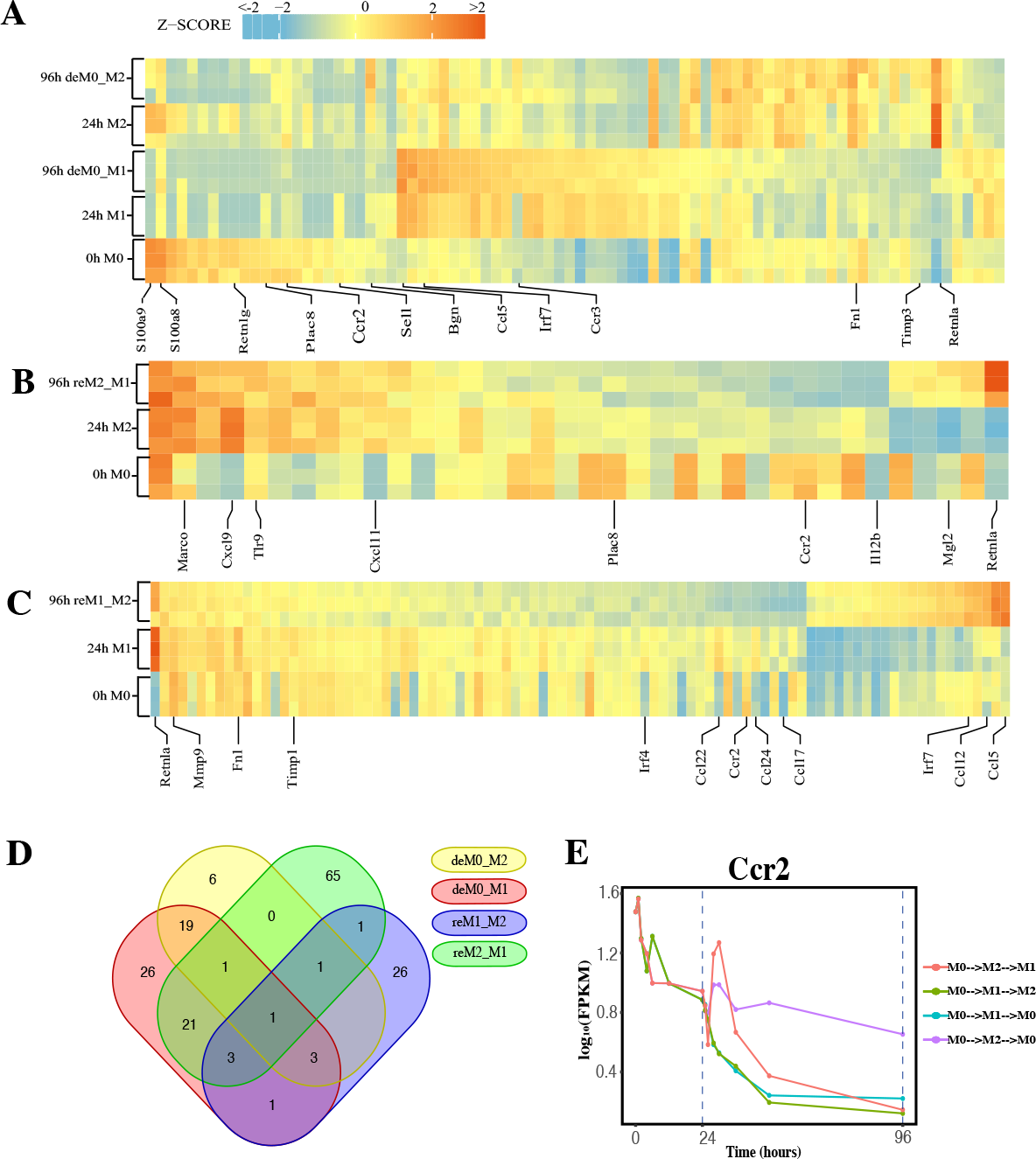
Identification of Hysteresis through differential expression analysis. (A–C). Expression pattern of significant hysteresis genes within **(A)** M0/M1/deM0_M1/M2/deM0_M2, **(B)** M2/reM2_M1, and **(C)** M1/reM1_M2. The FPKM were normalized to z-score scale. **(D)** Venn diagram illustrating the intersection of hysteresis genes. **(E)** Expression patterns of *Ccr2* over time in macrophage differentiations. The dashed lines indicate the key cell states at 24 h and 96 h time points. (M0->M2->M1: cell transition of M0 (at 0 h) to M2 (at 24 h) then to M1(96 h), M0->M1->M2: cell transition of M0 (at 0 h) to M1 (at 24 h) then to M2 (96 h), M0->M1->M0: cell transition of M0 (at 0 h) to M1 (at 24 h) then to M0 (96 h), M0->M2->M0: cell transition of M0 (at 0 h) to M2 (at 24 h) then to M0 (96 h))

Simultaneously, we applied identical criteria to screen for hysteresis genes between M1 and reM1_M2, and M2 and reM2_M1, illustrating two distinct types of hysteresis upon repolarization from M1 and M2. Our analysis identified 36 (Figure 2B, Supplementary Excel Sheet 3) and 93 (Figure 2C, Supplementary Excel Sheet 4) significantly differentially expressed hysteresis genes, respectively. These results underscore that macrophages in groups with a history of M1 polarization (deM0_M1 and reM2_M1) generated a greater number of hysteresis genes following either de-/repolarization.

Subsequently, in addition to the previously mentioned 24 common genes between deM0_M1 and deM0_M2, an additional 26 genes, including *Ccl5, Fn1*, and *Irf7*, were found to be common between deM0_M1 and reM2_M1 (Figure 2D). In contrast, the overlap of shared genes between the groups with a shared M2 polarization history (deM0_M2 and reM1_M2) was limited. This observation may stem from the relatively low number of hysteresis genes generated during M2 polarization. Remarkably, *Ccr2* was the only gene that showed hysteresis across all four groups. In our study, *Ccr2* expression peaked in the M0 state, significantly decreased after receiving M1 or M2 stimulation, and continued to decrease during the de-/repolarization process (Figure 2E). Notably, the blockade of the *Ccl2*–*Ccr2* axis has been associated with an elevated expression of M1 polarization-associated genes and a reduced expression of pro-M2 markers (35). *Ccr2* is also a known marker for immature monocytes, implying that M0 macrophages contain immature cells expressing *Ccr2*, and its expression decreases after maturation and is downregulated even after depolarization. However, few studies have investigated the role of *Ccr2* in macrophage hysteresis.

Among the common hysteresis genes with M1 history, *S100a8* and *S100a9*, which belong to the S100 family and function as Ca2+-binding heterodimers that modulate the inflammatory response of macrophages (36), were significantly downregulated. Existing evidence indicates that the impairment of *S100a9* expression during macrophage differentiation can cause an imbalance in inflammation and the wound healing processes (37). A significant reduction in the expression of *Plac8*, which can bind to *Cebpb* and control macrophage development and polarization (38), was observed in both de-polarized M0 and reM1_M2 groups. Similarly, as markers for M2 macrophages, the expression of *Retnla* and *Fn1* were downregulated in reM2_M1 cells compared to that in the M2 cells. In contrast, *Ccl5* and *Irf7* were the highly expressed hysteresis genes in deM0_M1 and reM2_M1 cells. These results suggest that the hysteresis gene produced by de-/repolarization may have an impact on the inflammatory response and developmental function of macrophages. Furthermore, the hysteresis induced by the M1 polarization history may be greater at the transcriptional level than that by M2 polarization history.

### Expression pattern of the hysteresis genes is similar to that of traditional macrophage marker genes

In addition to the established marker genes, we examined the expression patterns of other genes at the single-cell level. To achieve this, we leveraged the scRNA-seq data from the Liu project, encompassing unstimulated M0 cells at 0, 6, and 24 h, along with M1 cells with LPS & IFN-γ and M2 cells with IL-4 & IL-13 at 6 h and 24 h (Figure 1A) to validate their expression levels. After implementing the data pre-processing and quality control procedures, we amassed a total of 22,881 cells. We identified seven distinct cell subpopulations by appending their phenotype annotations (M0, M1, and M2). These subpopulations comprised M0 (0 h, 6 h, 24 h), M1 (6 h, 24 h), and M2 (6 h, 24 h) (Figure 3A).

**Figure 3:**
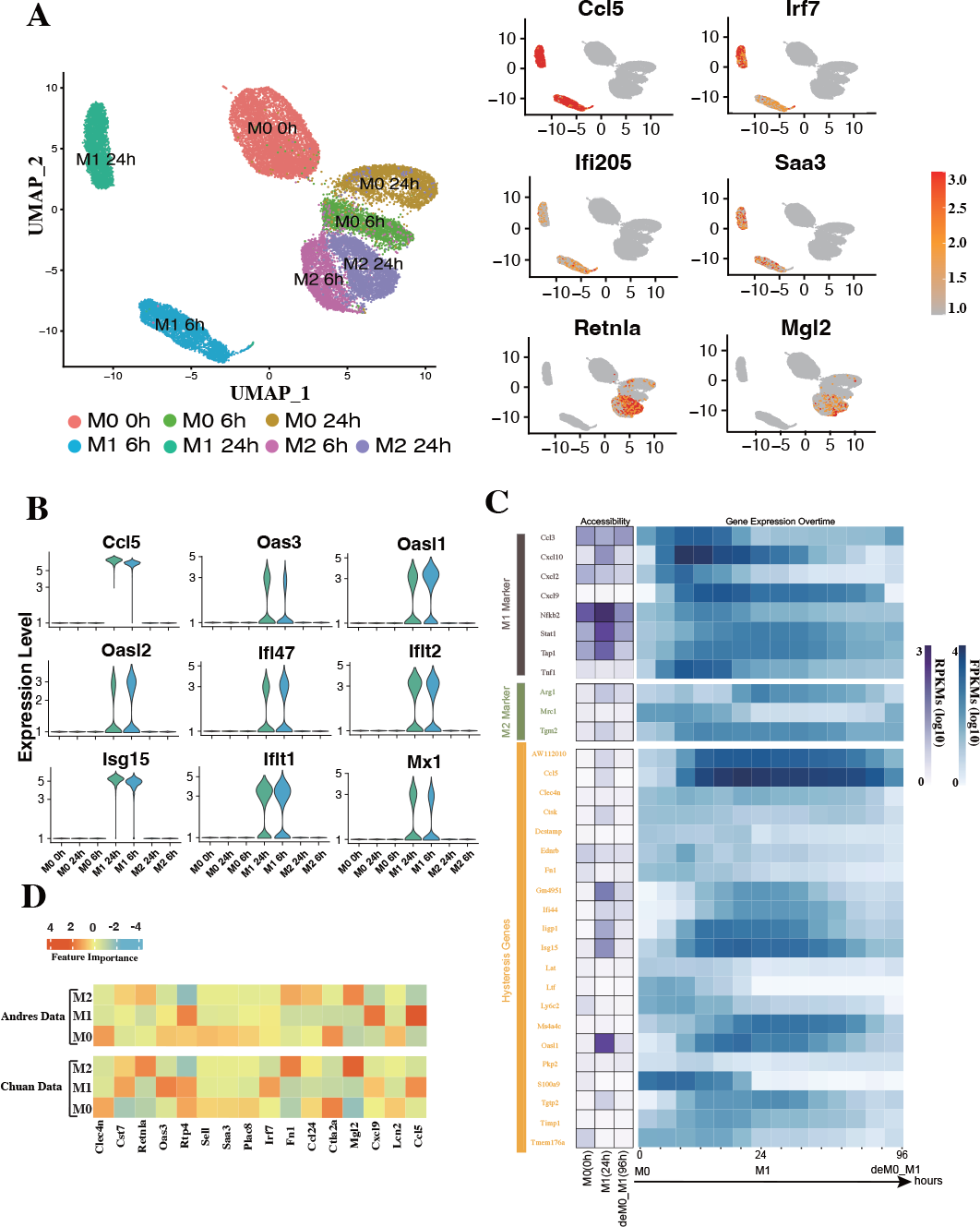
Hysteresis genes exhibit properties similar to traditional marker genes. **(A)** Umap visualization depicting gene expression patterns across different macrophage phenotypes at different stimulation time points. Hysteresis genes, such as *Ccl5, Irf7, Ifi205*, and *Saa3*, exhibit significant high expression in the M1 cell population, whereas *Retnla* and *Mgl2* show high expression in the M2 population. **(B)** Expression levels of hysteresis genes linked to M1 history. **(C)** Changes in chromatin accessibility at promoter regions and gene expression for selected M1 and M2 markers, along with hysteresis genes, during the M0->M1->deM0_M1 transition. **(D)** Scaled feature importance in logistic regression model for each of macrophage subtypes, evaluated using the Andres dataset (13) and Chuan dataset (14).

Subsequently, we evaluated the expression levels of hysteresis genes, particularly those with a history of M1 polarization. Some hysteresis genes (*Ccl5, Oas3, Oasl1, Oasl2, Ifi47, Ifit2, Isg15, Ifit1*, and *Mx1*) shared by both the deM0_M1 and reM2_M1 groups with M1 polarization history exhibited notable upregulation in the M1 phenotype (Figure 3B). Particularly, *Ccl5* and *Irf7* were consistently highly expressed in the M1 phenotype. Similarly, *Ifi205* (a conventional M1 marker) and *Saa3*, which are associated with M2 polarization, showed elevated expression within the M1 cluster (Figure 3A). The elevated expression of these genes within the M1 cell subset underscores the influence of the M1 polarization history on the comprehensive transcriptome of de-/repolarized macrophages. Interestingly, among the shared hysteresis genes in cells with an M2 polarization history (de_M0_M2 and reM1_M2), Less hysteresis genes (*Retnla* and *Mgl2*) exhibited a notably strong inclination toward high expression in the M2 cell population (Figure 3A). This aligns with our previous observation that the hysteresis effect is more pronounced in cells with an M1 polarization history than in those with an M2 polarization history.

We further analyzed whether the changes in the accessible chromatin regions of hysteresis genes were consistent with changes in gene expression levels by analyzing the ATAC-seq dataset (Figure 1A). After processing the data, we performed a comparative analysis of the changes in the chromatin accessibility of the promoter regions and RNA expression levels among the three states M0, M1, and deM0_M1. Although hysteresis genes exhibit sustained expression profiles, changes in chromatin accessibility in their promoter regions do not perfectly mirror the observed gene expression trends, akin to conventional marker genes. This suggests that the regulation of hysteresis gene expression may involve mechanisms beyond promoter region activation (Figure 3A).

Given the absence of apparent hysteresis in gene expression and chromatin accessibility among the marker genes, we investigated the significance of hysteresis genes in macrophage subtype classification. Employing a logistic regression classifier, we aimed to determine if hysteresis genes can be used to accurately classify macrophage phenotypes. We initially selected 16 hysteresis genes (*Ccl24, Retnla, Mgl2, Fn1, Cst7, Irf7, Plac8, Oas3, Lcn2, Ctla2a, Sell, Clec4n, Ccl5, Saa3, Rtp4*, and *Cxcl9*) as M0, M1, and M2 subtype features. Subsequently, a logistic regression model was trained using single-cell data from the Andres and Chuan research projects. Sixteen M1 and M2 markers (*Tlr4, Tlr2, Mhc2, Cd80, Cd86, Cxcl2, Ccl3, Stat1, Nfkb2, Tap1, Cxcl9, Cxcl10, Ccl8, Tlr1, Il1R, Tnf, Arg1, Mrc1, Nos2, Tgm2*, and *Cd163*), along with the PCA results, were used as positive controls. To establish a baseline, we randomly selected 16 genes from a single-cell dataset 100 times and averaged their performances. A grid search strategy was employed to optimize the hyperparameters for improved performance and mitigate overfitting concerns (refer to Methods). To achieve the optimal performance, we compared the prediction accuracy and F1 score (Table 1) derived from the confusion matrix (see Figure S1). The results indicate that hysteresis genes yielded high accuracy (0.87 in Andres data and 0.81 in Chuan data) and F1 scores (0.88 in Andres data and 0.79 in Chuan data) in comparison with randomly selected genes. However, this performance was surpassed by that of traditional M1/M2 macrophage markers (ACC: 0.89, 0.84; F1: 0.90, 0.83, respectively). Consistent feature importance trends (Figure 2D) across these genes in the model highlighted the reproducibility of classification outcomes; this verified that, while not achieving parity with marker genes, high accuracy underscored the unique expression profiles of hysteresis genes in M0, M1, and M2 macrophages.

**Table 1:** Result of the logistic regression model.

| Dataset | Random Select |  | Hysteresis Genes |  | Marker Genes |  | PCA |  |
| --- | --- | --- | --- | --- | --- | --- | --- | --- |
|  | ACC | F1 | ACC | F1 | ACC | F1 | ACC | F1 |
| Andres Pro-<br>ject (13) | 0.6735 | 0.6819 | **0.8784** | **0.8849** | 0.8987 | 0.9052 | 0.9977 | 0.9979 |
| Chuan Project<br>(14) | 0.7071 | 0.6934 | **0.8195** | **0.7954** | 0.8481 | 0.8372 | 0.9260 | 0.9244 |

These findings demonstrated that, similar to conventional macrophage markers, most hysteresis genes exhibit a propensity for elevated expression in either M1 or M2 subtypes. Notably, what distinguishes them is their ability to sustain expression patterns from the preceding state following de-/repolarization. This sustained expression/non-expression phenomenon could potentially exert an additional influence on functional modifications in de-/repolarized macrophages.

### Embedding of gene regulatory network revealed the functional impact of hysteresis genes on essential immunity, cell cycle, and cell migration in macrophages

To elucidate the roles of hysteresis genes, we performed functional enrichment analyses of the four distinct gene sets introduced in Section 3.1. GO and KEGG analyses (Figure 4A) revealed that hysteresis genes within the deM0_M1 and reM2_M1 groups exhibited pronounced enrichment in immune-related functions and pathways associated with M1 polarization, such as Type I interferon, NOD-like receptor, and TLR signaling pathways. Within the reM2_M1 group, we observed a notable enrichment of genes implicated in nuclear division and cell cycle functions. Conversely, cells with a history of M2 polarization showed limited enrichment, potentially due to the comparatively small gene pool. Notably, a shared enrichment of genes linked to cell chemotaxis was discernible across all four groups, implying a plausible connection between cellular migratory capacity and polarization history.

**Figure 4:**
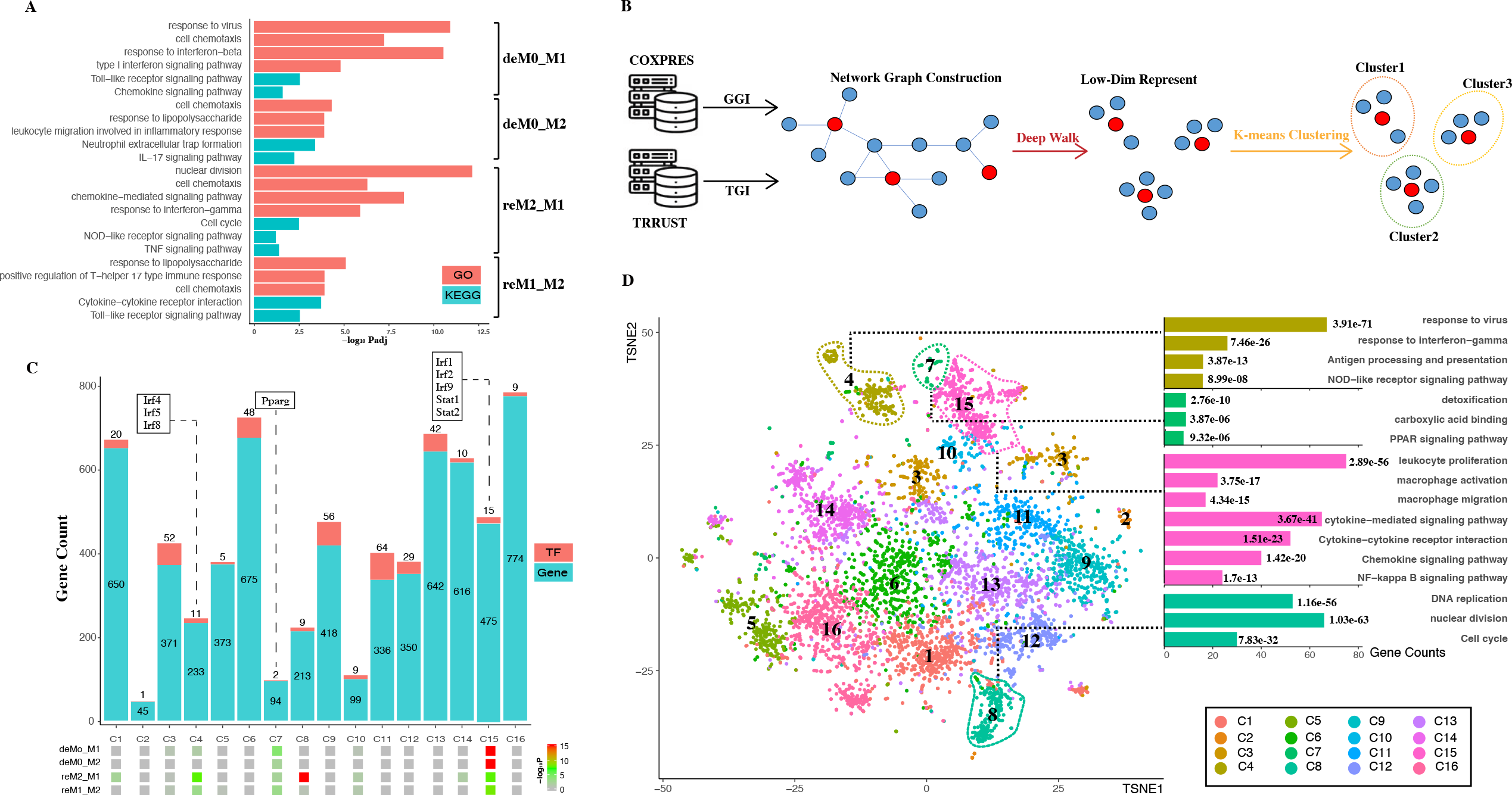
Hysteresis genes are enriched in gene clusters that influence multiple functions of macrophages. **(A)** GO and KEGG enrichment results measured in adjusted P-value in negative log-10 scale (-log_10_ Padj) for the four groups of hysteresis genes. **(B)** Schematic representation of graph embedding method and downstream analysis. A gene co-expression network was constructed by integrating gene–gene interaction (GGI) data from the COXPRES database and TF–gene interaction (TGI) data from the TRRUST database. Subsequently, the gene network was subjected to DeepWalk algorithm and K-means clustering to identify gene clusters for downstream analysis. **(C)** Enrichment levels of the four groups of hysteresis genes measured by P-value in negative log-10 scale (-log_10_ P) in each of the 16 gene clusters. The gene clusters are represented on the x-axis, and the number of genes and transcription factors in each cluster on the y-axis. **(D)** t-SNE (TSNE) plot reveals the distribution of all gene clusters within a reduced-dimensional space following graph embedding. The accompanying bar plot demonstrates significant GO terms and KEGG pathway enrichment outcomes corresponding to each gene cluster.

The genes were then embedded into a gene regulatory network (Figure 4B). This process harnessed the existing knowledge of macrophage TFs regulation and gene–gene co-expression from publicly available databases. Our approach initially involved constructing a network graph using both the gene co-expression matrix from the COXPRES database and TF gene regulation matrix from the TRRUST database. Subsequently, using DeepWalk, we projected this gene network graph into a lower-dimensional vector space to enable k-means clustering for predicting sub-gene regulatory clusters. Determining the number of subclusters entailed a joint consideration of the gene/TF counts and cluster functional annotations. Oversized (N > 800) or undersized (N < 30) gene clusters were meticulously avoided to ensure mutually exclusive cluster annotations. Finally, we chose 16 clusters (denoted as C1, C2, …, C16) for subsequent analyses. Among these 16 clusters (Figure 4C), all four hysteresis gene groups displayed significant enrichment in C15, which also incorporated critical TFs involved in type I interferon signaling (*Irf1, Irf2*, and *Irf9*) (39) and IFN response (*Stat1, Stat2*) (40). Although less pronounced than that in C15, all four hysteresis gene groups were enriched in C7, within Pparg that regulates macrophage intracellular lipid metabolism (41). Hysteresis genes linked to a shared M1 polarization history (deM0_M1 and reM2_M1) were notably enriched in C4, encompassing three additional members of the IRF family (*Irf4, Irf5*, and *Irf8*). Intriguingly, within all groups, significant enrichment at C8 was exclusively exhibited by the hysteresis genes of the reM2_M1 group.

Upon visualizing the network in a low-dimensional space (Figure 4D), we observed the spatial distribution of the 16 gene clusters using UMAP. Subsequently, we examined the GO annotations and KEGG pathways associated with each subcluster (Figure 4D). Our observations highlighted that C4, C7, and C15 are affiliated with antigen presentation, IFNγ response, Nod-like signaling/detoxification, PPAR signaling/macrophage activation, and NF-κB signaling, respectively. Notably, hysteresis genes from the reM2_M1 group were significantly enriched in C8, which was a cluster that exerted a substantial influence on DNA replication and the cell cycle, as indicated by the functional annotations.

Subsequently, we quantified transcriptomic disparities across the four groups (M0 vs. deM0_M1, M0 vs. deM0_M2, M1 vs. reM1_M2, and M2 vs. re_M2_M1). We used the Manhattan distance as a reference to gauge the overall gene expression differences. Treating each of the 16 subgene clusters as a distinct subset of overarching expression, we autonomously computed the expression disparities between the sub-clusters in de-/repolarized cells and the sub-cluster in the initial macrophage state. In our analysis, a sub-cluster was deemed to contribute to de-/repolarization-associated expression differences (i.e., hysteresis) if the Manhattan distance in a given comparison group exceeded the overall gene expression. Conversely, if a cluster’s Manhattan distance was low, it indicated a lack of or negative contribution to the expression differences. The results indicated that all four groups, C4, C7, C8, and C15, exhibited positive contributions to hysteresis (Figure S2). Comparatively, C8 showed substantial contributions to the overall differences compared to C15 in the deM0_M2 and reM2_M1 groups. However, the cumulative impact of minor expression differences among the genes, resulting from individual gene variations, significantly influenced overall expression disparities, despite the absence of conspicuous clustering of hysteresis genes. Collectively, these findings highlight the profound influence of differential hysteresis gene expression on essential immunity, cell cycle dynamics, and cell migration within macrophages.

### AP-1 motif is enriched in hysteresis chromatin region after M1 polarization and negatively correlated with the binding of CTCF

To investigate the underlying factors responsible for the hysteresis phenomenon between primary cells and de-/repolarized cells, we employed ImmunoNavigator online tools to predict enriched TFBS within the promoter regions of hysteresis-associated genes across the four groups (Table 2). Notable enrichment of TFBS was observed in the deM0_M1 and reM2_M1 groups, whereas no significant output was observed in the deM0_M2 and reM1_M2 groups, both of which had previously undergone M2 polarization. This aligned with our gene expression analysis, which indicated a lower expression of hysteresis-related genes compared with that of macrophages with an M1 polarization history. Furthermore, shared TFBS were identified between significant DEGs in the deM0_M1 and reM2_M1 groups. These TFBS include Irf1, a pivotal factor in M1 polarization (42); Irf2, known for inducing pro-inflammatory responses (43); and Stat1, a key mediator of M1 polarization induced by IFN-γ (44). To evaluate the specificity of this enrichment of hysteresis-associated genes only, we established a set of M1-specific DEGs using identical screening criteria. This yielded 365 M1-specific DEGs that were used as the control group (Supplementary Excel Sheet 5). In addition to Irf-related TFBS and Stat1, we identified enriched TFBS for NF-κB and Cebpa, which enable macrophages to modulate inflammatory responses via TLR4 stimulation. Our findings imply that IRFs-associated transcription factors and Stat1, rather than Nfkb1 or Cebpa, are potential upstream mediators of M1 hysteresis.

**Table 2:**
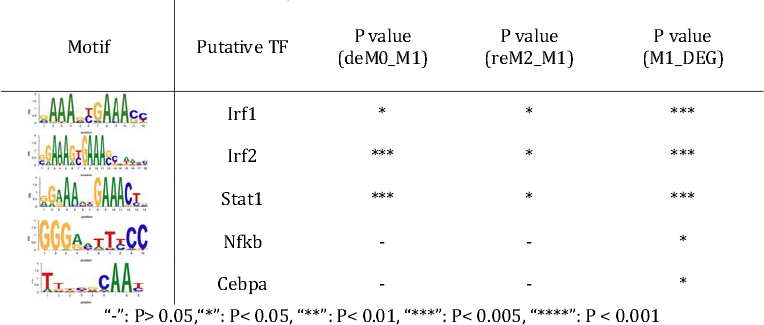
TFBS motifs enriched in promoter regions.

| Motif | Putative TF | P value<br>(deM0_M1) | P value<br>(reM2_M1) | P value<br>(M1_DEG) |
| --- | --- | --- | --- | --- |
|  | Irf1 | * | * | *** |
|  | Irf2 | *** | * | *** |
|  | Stat1 | *** | * | *** |
|  | Nfkb | - | - | * |
|  | Cebpa | - | - | * |

“-”: P>0.05, “\*”: P<0.05, “\*\*\*”: P<0.01, “\*\*\*\*”: P<0.005, “\*\*\*\*\*”: P<0.001

Next, we aimed to identify the cis-regulatory elements responsible for regulating the transcriptional identity of M1 hysteresis (Specifically, M0->M1->M0), focusing on chromatin accessibility. Liu et al. (9) involved three distinct ATAC-seq datasets corresponding to the M0, M1, and deM0_M1 states (Figure 1A). We aimed to detect changes in chromatin accessibility between primary macrophages (M0) and depolarized macrophages (deM0_M1) by analyzing these datasets. Our investigation of the accessible regions using the three ATAC-seq datasets consistently revealed strong signals for active cis-regulatory elements near the TSS regions (Figure 5A). However, we did not observe significant changes in chromatin accessibility following depolarization, which is consistent with the findings of the previous Liu project (9). Interestingly, both the overall ATAC-seq signal on TSS regions across the genome and on TSS regions of the hysteresis-related genes with significantly differential expression showed minimal differences in accessibility. Consequently, as discussed in Section 3.2, we propose that the expression of these hysteresis-related genes may not be primarily governed by direct binding events at the TSS regions, but rather by indirect regulation through distal regions.

**Figure 5:**
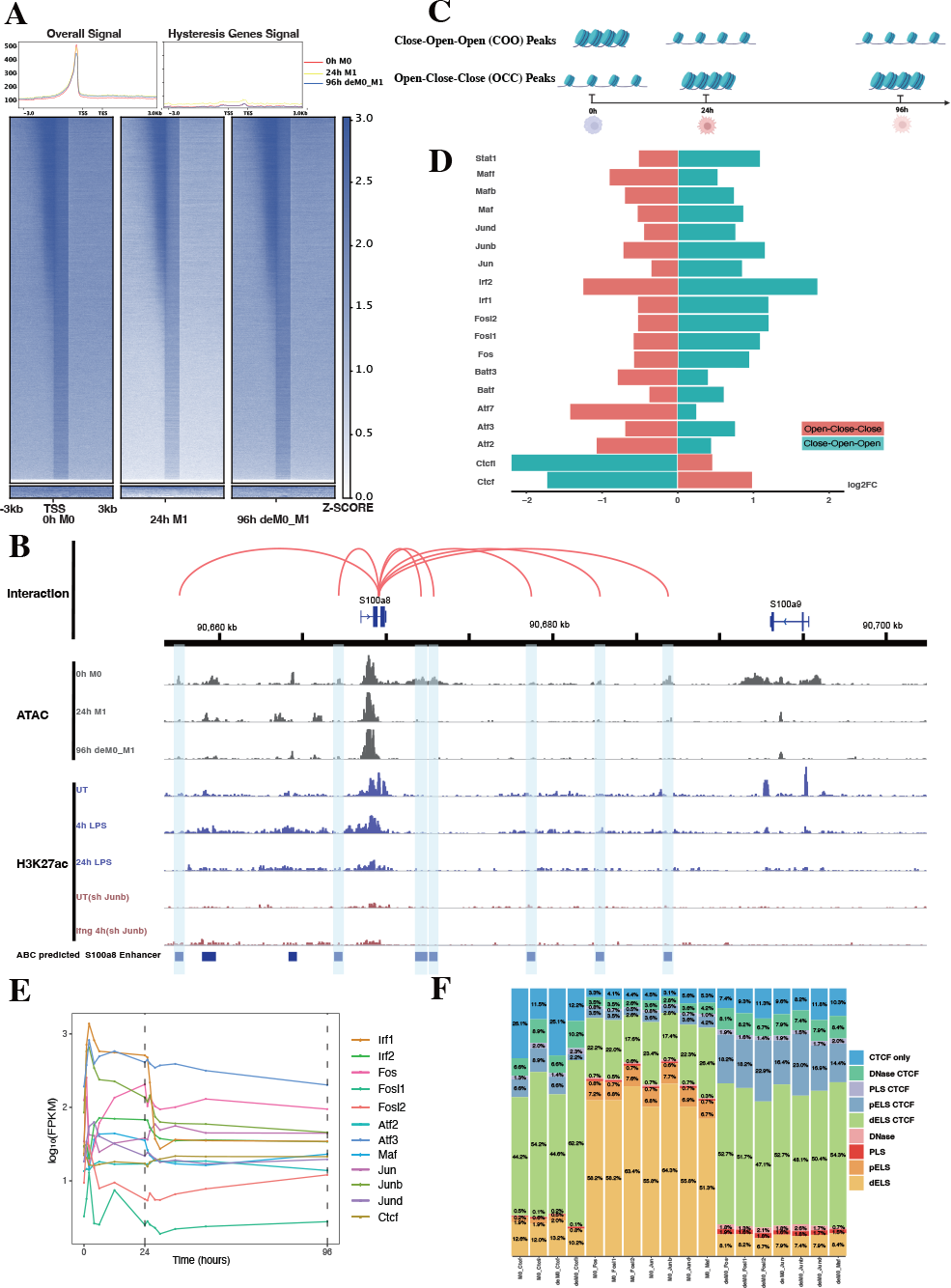
AP-1 and CTCF exert opposing regulatory effects on hysteresis. **(A)** ATAC signals in the upstream and downstream regions (3 kb) of TSSs (transcription start sites) of all genes and hysteresis genes were compared across the three cell states 0 h M0, 24 h M1, and 96 h deM0_M1. **(B)** Genome browser displays the potential gene–enhancer interactions within a 40-kb genomic region, including the genes *S1008* and *S100a9*. It shows the ATAC signal at 0 h, 24 h, and 96 h, as well as the H3K27ac signal in untreated (UT) macrophages, macrophages stimulated with LPS for 4 h and 24 h, untreated *Junb* knockdown macrophages, and *Junb* knockdown macrophages stimulated with IFN-_γ_ for 4 h. **(C)** Schematic representation of hysteresis region. The regions that exhibit low chromatin accessibility at 0 h and high chromatin accessibility at 24 h and 96 h are defined as the close-open-open regions. Vice versa, the regions that exhibit high chromatin accessibility at 0 h and low chromatin accessibility at 24 h and 96 h are defined as the open-close-close regions. **(D)** Log_2_(fold change) (validated by chi-square test) of the binding of AP-1, *Irf1/2, Stat1*, and *Ctcf/Ctcfl* in the open-close-close regions and close-open-open regions. **(E)** Gene expression levels of the AP-1 family and *Ctcf* at different time points. **(F)** Changes in *Ctcf* and AP-1 members binding sites within all the accessible regions of 0 h M0 macrophages and 96 h deM0 macrophages. (PLS (p): promoter-like signatures, ELS (e): enhancer-like signatures, DNase regions (d), and CTCF: CTCF-binding regions)

To validate our hypothesis, we employed the ABC method to predict enhancer regions within macrophages. Among the hysteresis-related genes, *S100a8* and *S100a9* were the most significant candidates (Figure S3). Notably, the promoter region of S100a9 showed a significant decrease in the peak signal from M0 to M1, which was maintained even after depolarization. This pattern was consistent with the ATAC and H3K27ac signal from 0 h to 24 h. In contrast, S100a8 did not exhibit any notable changes in its promoter region (Figure 5B). Intriguingly, we observed sustained closure of the predicted enhancer regions near the *S100a8* locus during the depolarization phase. This observation suggested a potential link between hysteresis and the cumulative impact of sustained opening and closure dynamics within gene-associated distal regions. However, upon further investigation of the peak variations of other hysteresis genes in this dataset, we discovered that this pattern could not be generalized to all genes.

Subsequently, we performed motif enrichment analysis on the open chromatin regions, discerning two distinct types of hysteresis peak regions (Figure 5C). The first type, termed open-close-close (OCC), demonstrated heightened chromatin accessibility in the M0 state, but diminished accessibility in both M1 and deM0_M1 states, suggesting sustained closure following depolarization. In contrast, the close-close-open (COO) region indicated the regions of persistent openness post-depolarization. Using the UniBind TFBS extraction tool, we derived the predicted motifs associated with each peak region. We then assessed the enrichment levels of these motifs in the OCC and COO regions relative to the overall peaks of high accessibility in the M0 and deM0_M1 states. Significantly enriched or depleted motifs within the OCC and COO regions were identified, and their log_2_(fold changes) compared to the initial state were computed (Figure 5D). Members of the IRFs (Irf1, Irf2), Stat1, and AP-1 (Maf, Mafb, Maff, Jun, Junb, Jund, Fos, Fosl1, Fosl2, Atf2, and Atf3) family were markedly depleted in the OCC regions, but significantly enriched in the COO regions. Interestingly, Ctcf and Ctcfl exhibited a reverse enrichment pattern, being prominently enriched in the OCC regions and depleted in the COO regions. The enrichment of Irf1/2 and Stat1 aligns with findings in this study on hysteresis gene promoters as well as the network analysis results implicating *Irf1,Irf2* and *Stat1* from C15 in hysteresis regulation. Collectively, these findings underscore the pivotal roles of the OCC and COO regions in the regulation of macrophage function, with AP-1, Stat1, IRFs, and CTCF occupying central regulatory positions. However, our investigation of the hysteresis gene promoter regions did not reveal the presence of AP-1 or CTCF, indicating that these transcription factors predominantly bind to distal regions for regulation, rather than to promoters. Notably, the expression levels of the AP-1 family and *Ctcf* showed no significant changes at 0 h and 96 h (Figure 5E), suggesting alterations in their binding activity, rather than their relative abundance during depolarization.

To gain deeper insights into the functional interplay between the AP-1 family and CTCF, we annotated mouse CTCF-bound cis-regulatory elements (cCREs) using SCREEN (45). This annotation categorizes cis-regulatory regions within the mouse genome into distinct signatures, namely promoter-like signatures (PLS), enhancer-like signatures (ELS), DNase regions, and CTCF-binding regions (CBRs), based on distinct histone marker signals. We focused on elucidating the changes in the binding patterns of AP-1 family motifs within all peak regions, comparing the 0 h M0 and 96 h de-polarized M0 states. We observed a notable shift in the motif-binding sites of members of the AP-1 family, including Junb, Fos, Fosl1, Fosl2, Jun, Jund, and Maf (Figure 5F). Specifically, in the M0 state, these motifs were predominantly enriched in regions devoid of Ctcf binding (non-Ctcf binding regions or non-CBR regions), with Junb accounting for 73.4% (Figure S4). However, in the deM0 state, these motifs were concentrated within the CBRs, with Junb occupancy of 88.1%. In comparison, the binding regions associated with Ctcf and Ctcfl exhibited no significant differences between the two states (Figure S5). These observations suggest that under depolarization conditions, AP-1 family members progressively bind to Ctcf-associated binding sites.

### ChIP-seq analysis revealed the role of AP-1 in hysteresis

To further dissect the regulatory interplay between the AP-1 family, CTCF, and a cohort of 75 M1 hysteresis genes (M0 -> M1 -> M0 group), we extracted PPI data from the STRING database. The network graph highlights the pivotal regulatory functions of some of AP-1 family members (*Jun, Junb, Jund, Fos, Fosl1, Fosl2, Maf, Atf3, Atf4*, and *Batf3*) and *Ctcf* within this interaction framework (Figure 6A). Notably, we also identified a cluster of ISGs in this network. These genes exhibited hysteresis in terms of promoter region accessibility (Figure 6B) and RNA expression levels (Figure 6C).

**Figure 6:**
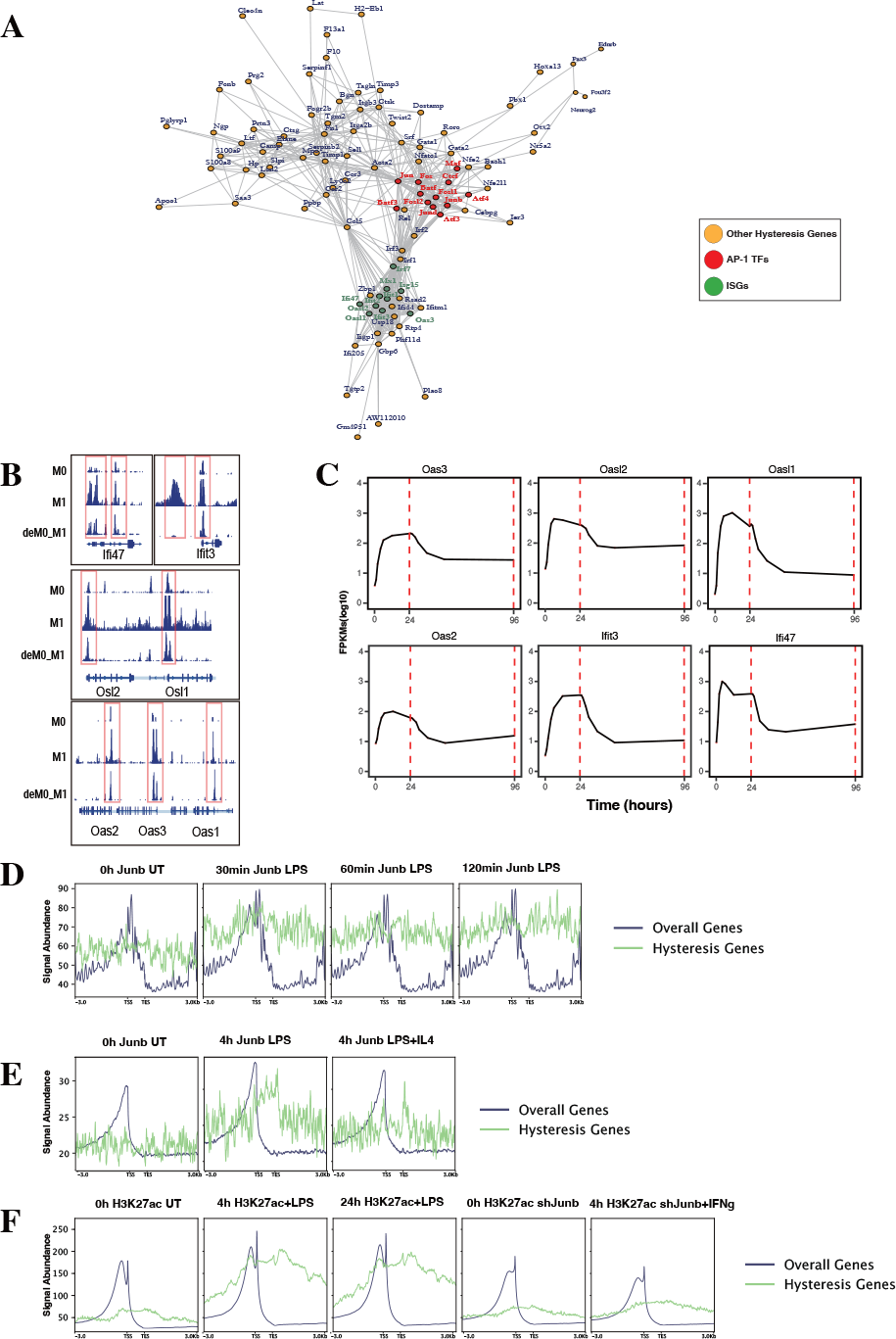
ChIP-seq analysis revealed the role of AP-1 in hysteresis. **(A)** Protein–protein interaction of hysteresis genes and AP-1/CTCF. **(B)** ATAC signal near promoter regions of ISGs across 0 h, 24 h and 96 h. **(C)** Change in the expression level of ISGs with time. **(D)** Binding of Junb to the promoter regions of all genes and hysteresis genes in BMDM changes upon LPS stimulation. **(E)** Binding of Junb to the promoter regions of all genes and hysteresis genes in BMDM changes upon 4 h LPS and 4 h LPS+IL4 treatment. **(F)** H3k27ac signal to the promoter regions of all genes and hysteresis genes in BMDM changes upon 4 h/24 h LPS treatment and 4 h IFN-_γ_+*Junb* knock down.

We compiled ChIP-seq datasets from BMDMs, encompassing Junb binding profiles from 0 to 2 h post LPS stimulation, Junb binding at 4 h post LPS stimulation, Junb binding following combined LPS and IL-4 stimulation, H3K27ac signals in BMDMs at 4 h and 24 h post LPS stimulation, and H3K27ac intensity in *Junb* knockdown BMDMs at 4 h post IFN-γ. Through rigorous ChIP-seq data analysis, we established that while Junb did not exhibit an evident enrichment tendency within the promoter regions of hysteresis genes, it did show an amplified binding signal in the promoter regions of hysteresis genes relative to the binding signals across all gene promoters after stimulation with LPS (Figure 6D). Interestingly, further validation using another set of ChIP-seq data revealed that the binding trend of Junb within the promoter regions of hysteresis genes after LPS stimulation remained unaltered by IL-4 (an M2 polarization cue) stimulation (Figure 6E). Moreover, during LPS stimulation, a notable enhancement in the H3K27ac signal was observed within the promoter region of hysteresis genes. Notably, in macrophages with *Junb* knockdown, the augmented H3K27ac intensity in the promoter region of hysteresis genes was no longer evident upon another type of M1 stimulation (IFN-γ) (Figure 6F). These findings collectively underscore the robust regulatory influence of *Junb* on hysteresis genes during M1 stimulation, which is not fully mediated by M2 stimulation. This dynamic interplay contributes to the emergence of the hysteresis phenomenon.

## Discussion

We sought to understand the molecular characteristics underlying macrophage de-/repolarization by integrating and re-analyzing large-scale publicly available datasets employing computational approaches. Liu et al. (9) demonstrated no significant global alteration in the transcriptome and chromatin accessibility of M0, M1, or M2 macrophages following de-/repolarization. Our findings reaffirmed this observation. However, our investigation specifically concentrated on a subset of genes with limited, yet notable, differential expression after de-/repolarization (designated hysteresis genes), as well as regions exhibiting substantial differences in chromatin accessibility. Our observations encompass the following facets: 1. A number of genes maintained their expression patterns during polarized state (persistently high or persistently low expression). This phenomenon is termed “macrophage hysteresis”, and the genes manifesting this behavior are categorized as hysteresis genes. Notably, the persistence of M1 polarization gave rise to a more pronounced form of hysteresis than that induced by M2 polarization. 2. These hysteresis genes exhibited expression trends analogous to those of conventional macrophage marker genes and played a pivotal role in macrophage phenotype classification. 3. The functional attributes of these hysteresis genes were interconnected with fundamental immunological functions, such as inflammatory responses, cell cycle, and cell migration within macrophages. 4. In macrophages undergoing M0 -> M1 -> M0 polarization, IRFs and Stat1 were enriched in the promoter regions of hysteresis genes. Conversely, AP-1 and CTCF bounded to mutually distinct distal regions (the COO and OCC regions, respectively). Collectively, these factors orchestrate regulating macrophage hysteresis (Figure 7) and trigger the cellular functional heterogeneity of macrophages.

**Figure 7:**
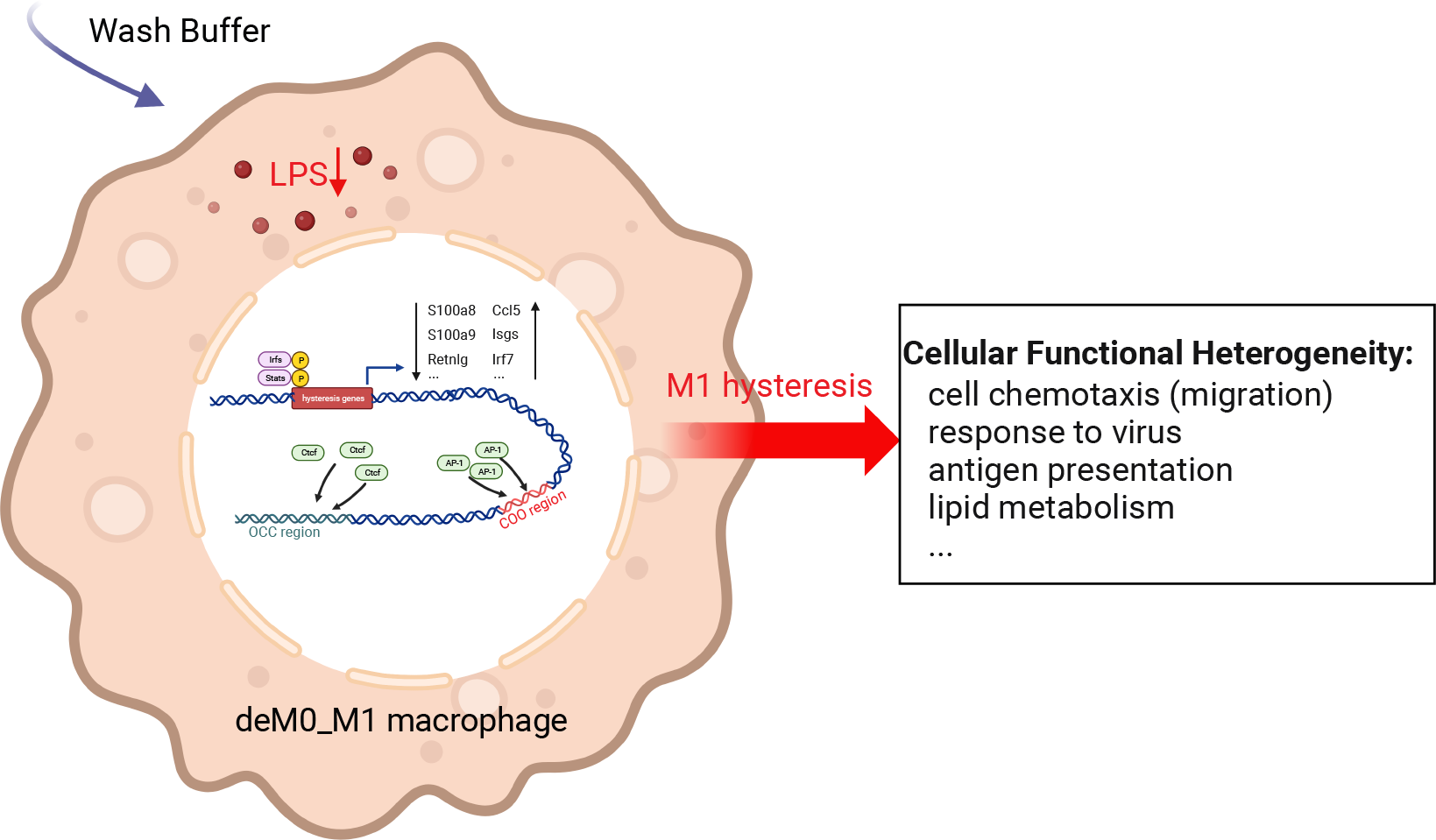
Potential regulatory mechanisms underlying M1 hysteresis. Upon withdrawal of the M1 stimulus (LPS) with wash buffer in polarized M1 macrophages, IRFs and STAT exhibit binding affinity towards the promoter regions of hysteresis genes, whereas AP-1 and *Ctcf* engage with the COO (Close-Open-Open) and OCC (Open-Close-Close) regions of chromatin, respectively. The conjoint orchestration of regulatory elements spanning both promoter and distal regions underpins the sustained expression profile of hysteresis genes. This transcriptional divergence subsequently reverberates across functionalities encompassing cell migration, antiviral response, antigen presentation, and lipid metabolism.

Given the opposing enrichment patterns, it is conceivable that AP-1 and CTCF may exert antagonistic effects on macrophage hysteresis by competing for binding sites, thereby inducing extensive alterations in the regulatory landscape. Although the relationship between AP-1 and CTCF remains unclear, in the human A549 cell line, CTCF depletion occurred at the hotspots enriched with AP-1 motifs (46). Within the context of inflammatory responses, the TLR4-NOS1-AP1 signaling axis governs macrophage polarization by facilitating AP-1 interactions and inducing IL-12/IL-23 (47). The AP-1 family including Fos and Jun (48) can co-modulate with NF-κB to regulate macrophage inflammatory responses (49). Furthermore, CTCF depletion attenuates acute inflammatory responses in induced macrophages (50). CTCF also controls proinflammatory gene expression through TLR signaling in macrophages (51). In addition to macrophages, through the regulation of the 3D enhancer network, CTCF participates in the inflammatory response of bone marrow-derived dendritic cells. Thus, both AP-1 and CTCF regulate inflammatory responses in cells. We identified a potential antagonistic relationship between AP-1 and CTCF; however, the precise mechanistic underpinnings require further investigation.

In our study, inducible ISGs exhibited hysteresis phenomena in response to M1 stimulation and displayed a robust correlation with AP-1 and CTCF within the PPI network of hysteresis genes. The downregulation of these type-I IFN genes serves as a protective mechanism against excessive lipoprotein uptake in human and murine foamy macrophages (52). Notably, another significant hysteresis gene, *Ccl5*, demonstrated tight PPIs with these ISGs. Elevated *Ccl5* expression has been linked to epigenetic modifications induced by AP-1 in adipose tissue macrophages (ATMs) of mice fed a high-fat diet (53). Furthermore, in the context of obesity, *S100a8/S100a9*, which are significant hysteresis genes recognized as damage-associated molecular patterns (DAMPs), are presumed to trigger ATM-mediated immune responses (54–56). Interestingly, our findings suggested that the hysteresis of *S100a8/S100a9* is controlled by the accessibility of the promoter and enhancer regions. This evidence suggests that, apart from the aforementioned cellular and immune functions, the differential expression of hysteresis genes is potentially linked to macrophage lipid metabolism. This intriguing avenue is a potential direction for future in-depth studies.

Polarization is thought to have little impact on macrophage memory, as traditional macrophage marker genes showed minimal differences in transcriptome and chromatin accessibility (9). However, we found that even subtle differences in expression at the cellular level can lead to functional changes in macrophages. Functional changes are likely to be mediated through epigenetic reprogramming, which is critical for adaptive immunity. For instance, a history of diet-induced obesity permanently alters chromatin accessibility (53).

However, several important points need to be considered. Firstly, our study has been built upon the analysis of data from multiple distinct projects, and as such, we were unable to completely control the batch effect due to various experimental conditions (dose of M1/M2 stimulus, etc.). Secondly, due to limitations in available data, we were unable to dissect the mechanisms underlying M2 hysteresis to the same extent as we did for M1 hysteresis. Furthermore, we could only consider the 96-hour as the end point of macrophage de-/repolarization, the hysteresis beyond the 96-hour stimulation period has not been discussed. Lastly, all the data were derived from mouse BMDM in vitro conditions. Therefore, the results may not directly correspond to tissue-resident macrophages in vivo conditions.

Taken all together, our approach allows the establishment of a multi-omics profile of macrophage differentiation and promotes our understanding of cellular hysteresis, which contributes to the future development of novel macrophage repolarization therapies. Hysteresis has been hypothesized in other immune-related cells, and further efforts are needed to understand the general and cell-type-specific hysteresis mechanisms.

## Supporting information

Supplementary Excel Sheet 1

Supplementary Excel Sheet 2

Supplementary Excel Sheet 3

Supplementary Excel Sheet 4

Supplementary Excel Sheet 5

## Acknowledgements

Computational resources were provided by the supercomputer system SHIROKANE at the Human Genome Center, Institute of Medical Science, University of Tokyo.

## Author contributions

YZ, YK, and KN conceived and designed the study. YZ, WY, and ML performed the analyses. YZ, YK, ML, SJP, and KN drafted the manuscript. All the authors have read and approved the final version of the manuscript.

## Materials and Methods

### Data preparation

The time-course bulk RNA-seq, single-cell RNA-seq (scRNA-seq), and bulk ATAC-seq data used in this study were sourced from the Liu Project (9) (GSE158094). The scRNA-seq data used to train logistic regression classifier were obtained from the Andres project (GSE117176) (13) and the Chuan project (GSE117176) (14). The ChIP-seq data employed for enhancer prediction were acquired from the ENCODE project (GSM1000074). The ChIP-seq data used to validate the role of AP-1 were obtained from the GEO database (GSE38377, GSE36099, GSE36104, GSE84519, and GSE84520). All the data in this study were obtained from mouse bone marrow-derived macrophages (BMDM) in the in vitro condition.

### NGS data processing

#### Bulk RNA-seq data analysis

The log10 normalized fragments per kilobase million (FPKM) mRNA expression matrices from the primary (M0), polarized (M1 and M2), de-polarized (deM0_M1, deM0_M2), and re-polarized (reM1_M2 and reM2_M1) macrophages were directly downloaded from GEO and used for downstream analysis. Differentially expressed genes (DEGs) were selected based on average difference of log10(FPKM) > 0.5, and P values < 0.05 between M0/M1/M2 (control) and 96 h de-polarized M0/re-polarized M1/M2, respectively. P-values were calculated using Student’s t-test.

### Single-cell RNA-seq analysis

Seurat (version 4.2.0 (15)) was used for preprocessing and downstream analysis of the macrophage single-cell RNA-seq (scRNA-seq) data from the Liu project. First, the macrophage phenotype annotation provided in the original article was used to differentiate single-cell macrophages. Thereafter, the selected cells with 200–2500 detected genes, > 200 nUMIs, and < 10% mitochondria genes were used for quality control and performed downstream analyses. Subsequently, 22752 macrophage cells with 4085 gene expressions were extracted and normalized using the standard Seurat protocol. The top 3000 variable features were selected, and the cells were clustered and visualized in the uniform manifold approximation and projection (UMAP) plot. Canek (16) was used to remove batch effects from the single-cell data. Raw counting matrices from the Andres and Chuan projects were used directly for logistic regression without any additional processing.

### Bulk ChIP-seq and ATAC-seq analyses

The raw ChIP-seq/ATAC-seq SRR files were downloaded from Sequence Read Archive (SRA). After converting to FASTQ format, reads were checked and trimmed by FastQC (17) and Trim Galore (18). Read alignments were performed against the GRCm38/mm10 reference mouse genome using Bowtie2 (version 2.4.5) (19), with the default settings. BAM (*. bam) files were generated and merged using samtools (1.3.1) (20). Low-quality alignments and mitochondrial reads were detected and removed using the Sambamba software (0.7.1 (21)); peaks were called by MACS2 (2.1.1) (22). Plotting of ChIPseq/ATAC-seq signal intensities in the transcription start site (TSS) region, conversion of files to BigWig format, and RPKM normalization were performed using deep tools (23) for subsequent analysis. The peaks in the BED files of the genome were visualized using IGV (24).

### Logistic regression on scRNA-seq data

Logistic regression classification was performed using the scikit-learn (25) python package. Raw scRNA-seq counts from the Andres and Chuan projects were used as inputs. Eighty percent of the samples was selected as training data and the remaining 20 percent was selected as testing data. An OVR (one vs. rest) strategy was used to identify the subtypes of macrophages. The hyperparameters were selected using the GridSearchCV strategy to minimize the loss function and achieve the best performance. The cell-type label provided by the original research was considered the ground truth to calculate the accuracy and F1 score. Finally, the feature genes and their importance were selected using the trained classifier.

### Network construction, Graph embedding, and Clustering analysis

To construct the ground truth of the TF-Gene interaction network in mice, gene co-expression data were collected from COXPRESdb (26), and the coexpression data were filtered based on the co-expression score provided by the database. Specifically, gene pairs with co-expression scores > 5 were retained. Confidence in the TF-Gene regulation data was acquired from the TRRUST database (27). Subsequently, the information from the two data sources was merged and a subnetwork of genes, which only existed in our time-course bulk RNA-seq dataset from the Liu project, was selected to construct a TF–gene interaction network.

In our network, 6776 gene nodes within 382 transcription factors (TFs) were identified. If two nodes had either a co-expression or a regulatory relationship, an undirected edge connecting the nodes was added to the graph. Given the undirected graph G = [V, E], where V represents all selected genes and E represents all co-expression or TF regulation relationships; the weight of E was not considered. Graph embedding analysis was performed using DeepWalk (28). For an unweighted graph, by performing a random walk method, DeepWalk could resemble the node sequences in the graph, which are then input into Word2Vec to perform the graph embedding. OpenNE (https://github.com/thunlp/OpenNE), which is an open-source Python package, was used to apply the DeepWalk embedding method. The K-means clustering method was directly applied to the embedding matrix, where the K value was chosen based on the number of genes in each cluster, with 16 selected as the final K value. Subsequently, Gene Ontology (GO) and Kyoto Encyclopedia of Genes and Genomes (KEGG) enrichment analyses were performed using ClusterProfiler (29) for cluster function annotation. GO terms and KEGG pathways with corrected P-values < 0.05 were selected as significant annotations. The enrichment of the hysteresis genes in each cluster was calculated using Fisher’s Exact Test.

### Transcription factor binding site (TFBS), enhancer prediction, and hysteresis peaks identification

UniBind (30) was used to predict the potential TFBS among all ATAC peaks. The selected DEGs were uploaded to the ImmunoNavigator (31) online tool to predict potential TFs regulating gene expression on promoter regions. The enhancer-gene interactions were predicted using the activity-by-contact (ABC) method (32), following documentation from the ABC repository in GitHub and setting up the conda (Anaconda, 2020) environments and suggested packages. Subsequently, both the processed ATAC-seq peaks and H3K27ac peak files were input. Candidate Region function with the recommended parameters (nStrongestPeaks 150000 and peakExtendFromSummit 250) was used. Enhancer activity was quantified using the ABC run.neighborhoods function with the mm10 gene as a reference. Finally, the ABC scores of enhancer-gene interactions were computed with the ABC prediction function setting –scale hic_using_powerlaw and a threshold of 0.2.

The following two types of hysteresis regions were defined: the open-close-close (OCC) region, representing the regions that are highly accessible at M0(0h), but significantly less accessible at M1(24h) and deM0_M1(96h); and the close-open-open (COO) region, representing the regions that are significantly less accessible at M0(0h), but significantly accessible at M1(24h)/deM0_M1(96h). Significant peaks that overlapped these regions were selected as OCC/COO peaks for downstream analysis using BedTools (v2.30.0) (33). The significance of the results was determined using the chi-square test.

### Other bioinformatics analysis and statistical analysis

The CTCF-binding annotation was downloaded from ENCODE (https://www.encodeproject.org/data/annotations/). Protein–protein interaction (PPI) analysis was performed based on the data from the STRING database (34). P<0.05 was considered statistically significant, and the other p-value conclusions were as follows: “-”: P> 0.05, “*”: P< 0.05, “**”: P< 0.01, “***”: P< 0.005, and “****”: P< 0.001.

**Figure S1:**
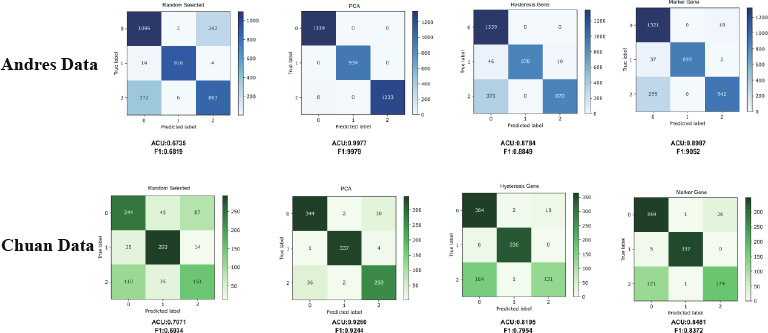
Confusion matrix of logistic regression results across each group (Random, PCA, hysteresis genes, and marker genes) within the Andres and Chuan datasets. The x-axis represents the predicted labels and y-axis represents the true labels. The numbers in the matrix represent cell counts.

**Figure S2:**
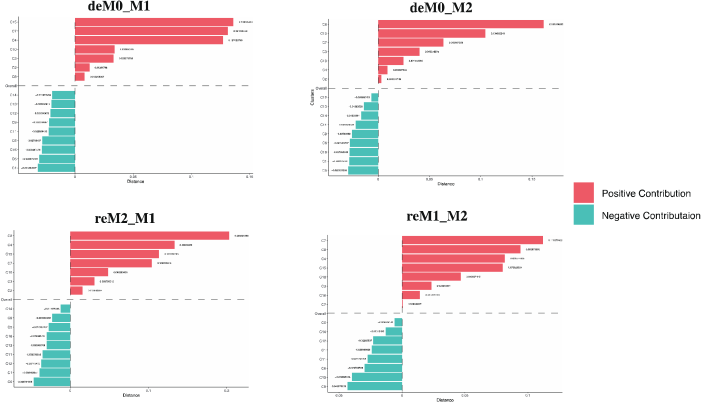
The four graphs represent the contribution of each gene cluster to the differences in the expression between the starting and endpoints of macrophage polarization, as measured by the Manhattan distance. Each graph corresponds to a specific polarization-de-/repolarization trajectory. The y-axis represents the gene clusters and x-axis represents the contribution of each cluster to the overall difference.

**Figure S3:**
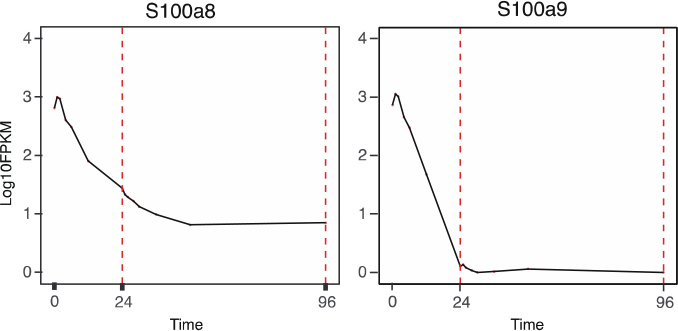
Gene expression levels of *S100a8* and *S100a9* were measured at different time points during polarization/depolarization. The x-axis represents time and y-axis represents gene expression levels (log_10_FPKM).

**Figure S4:**
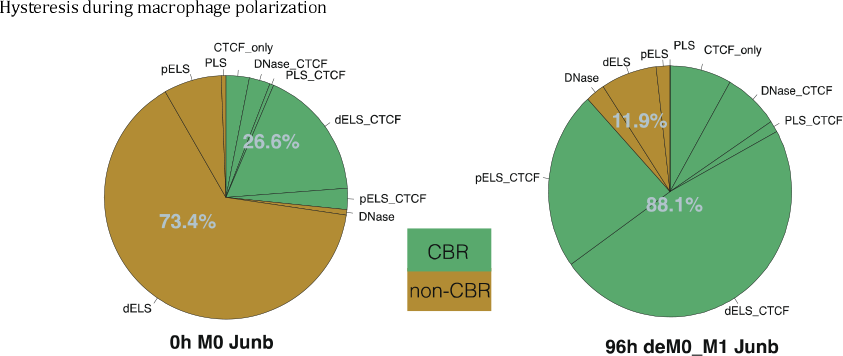
Changes in Junb motif-binding sites within all ATAC-seq accessible regions of 0 h M0 macrophages and 96 h deM0 macrophages.

**Figure S5:**
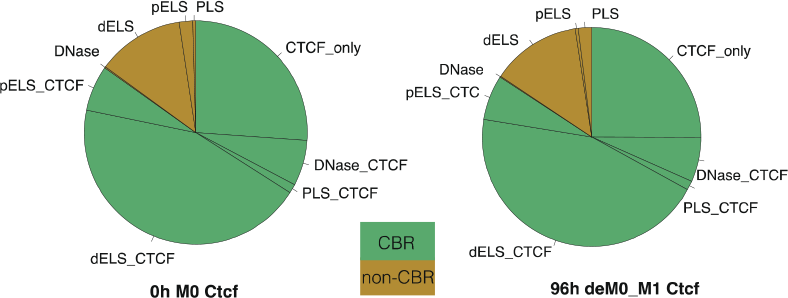
Changes in Junb motif-binding sites within all ATAC-seq accessible regions of 0 h M0 macrophages and 96 h deM0 macrophages.

